# Physiological responses to a changing winter climate in an early spring-breeding amphibian

**DOI:** 10.1101/2024.01.03.574103

**Authors:** Robin Schmidt, Cecile Zummach, Noa Sinai, Joana Sabino-Pinto, Sven Künzel, Kathrin H. Dausmann, Katharina Ruthsatz

**Author notes:** Robin Schmidt and Katharina Ruthsatz contributed equally. Corresponding author: **Katharina Ruthsatz**; ORCID: 0000-0002-3273-2826. Current address: Zoological Institute, Technische Universität Braunschweig, Mendelssohnstraße 4, 38106 Braunschweig, Germany.

## Abstract

Climate change is swiftly altering environmental winter conditions, leading to significant ecological impacts such as phenological shifts in many species. As a result, animals might face physiological mismatches due to longer or earlier activity periods and are at risk of being exposed to late spring freezes. Our study points for the first time to the complex physiological challenges that amphibians face as a result of changing thermal conditions due to winter climate change. We investigated the physiological responses to a period of warmer winter days and sudden spring freeze in the common toad (*Bufo bufo*) by acclimating them to 4°C or 8°C for 48 h or exposing them to 4°C or −2°C for 6 h, respectively. We assessed the daily energy demands, determined body condition and cold tolerance, explored the molecular responses to freezing through hepatic tissue transcriptome analysis, and measured blood glucose levels. Toads acclimated to higher temperatures showed a higher daily energy expenditure and a reduced cold tolerance suggesting faster depletion of energy stores and the loss of winter acclimation during warmer winters. Blood sugar levels were higher in frozen toads indicating the mobilization of cryoprotective glucose with freezing which was further supported by changed patterns in proteins related to glucose metabolism. Overall, our results emphasize that increased thermal variability incurs physiological costs that may reduce energy reserves and thus affect amphibian health and survival. This might pose a serious threat to breeding adults and may have subsequent effects at the population level.

## 1. Introduction

Understanding the responses and adaptability of wildlife to climate change is still a major challenge in ecological research. To date, most of the research exploring the impacts of climate change on animals has focused on effects of extreme heat events as well as on how animals might cope with environmental temperatures close to their upper thermal limits (e.g., 1;2;3). However, in many northern temperate ecosystems, shifts in temperature and precipitation patterns have been, and are expected to be, more pronounced during winter compared to summer (4;5). *Winter climate change* is driving complex changes in both thermal means and temperature extremes with a generally increasing trend (rev. in 6). Particularly at high latitudes, climate is warming significantly faster in winter than in summer (7;8). Furthermore, some regions are experiencing unusual cold spells or extreme weather events as a result of climate-induced changing circulation patterns, making winters more variable in addition to warmer (9).

A common ecological consequence of winter climate change across diverse taxa is phenological shifts in critical life-history events (10;11;12;13). These include an accelerated onset of the growing season for many plants (14), an advanced return of in migratory birds (15), the anticipated emergence of many dormant insect or mammal species (16;17), as well as an earlier initiation of the breeding season (18). Among all taxa, amphibians have been shown to reveal the strongest average shift towards earlier breeding (19), with more than 60% of all populations studied doing so (20). However, the magnitude of phenological response is known to be species- and population-specific, and to vary with latitude (19;21).

With a shifted breeding phenology induced by winter warming, adult amphibians are likely experiencing an extended or earlier annual activity period. Since temperate amphibians rely on internal energy reserves (i.e., the fat body) during overwintering, warmer winter temperatures as well as extended or earlier activity periods might increase metabolic costs and drain such energy reserves (22;23) with consequences for body condition and survival (24;25). In addition to warming, climate change will also increase the occurrence of extreme weather events such as sudden frost in spring known as *false springs* (26;27), thereby exposing amphibians to the increased risk of cold stress. Despite an increased risk of being frozen to death or frost damage, experiencing cold stress after the reproductive season has begun is physiologically costly (28). This cold stress has many adverse effects including reduced energy reserves, comprised immune system functioning, and decreased fecundity (rev. in 29), and might therefore pose a serious threat to breeding amphibians.

Amphibians have developed several responses to cope with cold stress such as the selection of microhabitats with more favorable temperatures to mitigate cold exposure (30;31) or physiological adaptations to endure cold conditions (32;33). If behavioral thermoregulation is constrained, such as through the occurrence of sudden frost, freeze protection or tolerance through the accumulation of cryoprotectants, such as plasma glucose derived from liver glycogen catabolism, glycerol, or urea, might contribute to withstanding extreme cold exposure (34;35;36). However, if energy reserves are drained due to warmer winter temperatures as well as extended or earlier activity periods, the ability to successfully implement an effective response to cold stress might be impaired (6). Consequently, increases in mean temperature could potentially increase the susceptibility to extreme weather events such as sudden frost. Nonetheless, the physiological challenges and potential impacts of changing winter climate on amphibians are still not well understood.

In this study, we investigated the physiological responses of early spring-breeding temperate amphibians to changing thermal conditions resulting from winter climate change, using the common toad (*Bufo bufo*) as a model species. Specifically, we aimed to determine metabolic costs resulting from higher winter temperatures combined with exploration of cryoprotective mechanisms in response to sudden (night) freeze. To simulate warmer winter days and spring freeze, toads were acclimated to 4°C or 8°C for 48 h (i.e., experiment 1) or exposed to either 4°C or −2°C for 6 h (i.e., experiment 2), respectively. We measured the standard metabolic rate to assess daily energy demands and determined body condition, cold tolerance as well as glucocorticoid hormone levels as a proxy for physiological stress in toads acclimated to different winter temperatures. To explore the molecular responses to freezing, we used a transcriptome-level investigation of hepatic tissue, and measured blood glucose level. We predicted that toads acclimated to warmer winter temperature would experience reduced cold tolerance, higher levels of physiological stress, and increased metabolic rate, resulting in higher mass loss and depletion of energy budgets compared to the control group. We further predicted that toads facing false springs would respond with the mobilization of cryoprotective glucose and reveal differentially regulated hepatic pathways related to cryoprotectant mobilization and stress tolerance.

## 2. Materials and Methods

We ran two experiments to investigate the physiological responses to warmer winter days (i.e., *experiment 1: Warmer winter scenario*) and sudden spring freeze (i.e., *experiment 2: Spring freeze scenario*) in the widespread anuran *B. bufo*. This early spring-breeding amphibian was selected as a model species in this study because its spring migratory and spawning behaviour directly exposes it to changes in temperature caused by winter climate change, and its reproductive success can be strongly influenced by these changes (37).

### 2.1 Field work and animal collection

The fieldwork was carried out at the locality Kleiwiesen (52.328 N, 10.582 E) near Braunschweig, Germany, after the migration of *B. bufo* had started on March 17^th^ and 18^th^, 2022 (Table S1). This site sustains a large population of *B. bufo*, which breed in a shallow part of one pond, partly covered with dense reeds. In total, we collected 40 toads (i.e., n_experiment1_=30; n_experiment2_=10) by hand wearing nitrile gloves and kept them individually in small plastic boxes containing wet tissue to avoid dehydration (Table S1, S2). Special care was taken to capture only male toads on their way to the breeding pond to avoid any impact on the population, considering that many female toads typically have just one opportunity to spawn in their lifetime (38). Mean ambient temperature during the toad collection was 3.72 °C (Table S1). Therefore, we considered 4 °C as temperature threshold for the start of the migratory activity in male toads (82 but see: 83) and used this temperature as control temperature in both experiments. After collection, the toads were transported to the laboratory on the day of collection, where their snout-vent-length (in cm; SVL) and body mass (in g) were measured. SVL was measured with a caliper (to the closest 0.05 cm) and body mass was determined with an electronic balance (to the nearest 0.01 g; Sartorius A200 S, Göttingen, Germany) (Table S1). As digestion is accompanied by an increase in metabolic rate, animals were fasted for 48h (i.e., during the thermal acclimation period) before the physiological measurements (39). This fasting mimics natural conditions as during the migration to the breeding ponds, spring breeding amphibians rely on their internal fat storages (i.e., fat bodies; 22).

### 2.2 Experiment 1: Warmer winter scenario

#### 2.2.1 Animal husbandry and acclimation conditions

To test how a period of warmer winter days affects the physiology of *B. bufo*, 30 animals were randomly split into two groups and acclimated to 4°C and 8°C for 48 hours, respectively, using two different refrigerators. Acclimation temperatures were chosen in order to simulate thermal conditions during early spring (i.e., 4 °C) and warmer conditions (i.e., 8 °C) as predicted under changing global climate following the approach of Podhajský & Gvoždík (84). Refrigerators were set to the respective acclimation temperature and equipped with additional digital thermometers (TFA LT-102, Conrad electronics, Hirschau, Germany; precision: ± 0.5 °C). Animals were housed individually in plastic boxes (10 × 20 × 15 cm) with moist tissue to prevent dehydration. Tissues were changed daily, and new water was added to keep the boxes clean.

#### 2.2.2 Physiological measurements

After the 48-hour acclimation period, the following physiological parameters were determined: body condition, water-borne corticosterone (CORT) release rates, standard metabolic rate (SMR), daily energy expenditure (DEE), and cold tolerance (Table S1). After the physiological measurements, all 30 toads were kept at 4°C for an additional 48 h and fed medium-sized house crickets (*Acheta domesticus*) *ad libitum* to allow them to recover and feed before we returned them to the collection site.

##### Body condition

Body condition was determined directly after collection as well as after 48h (i.e., acclimation duration) to examine if the body condition changed during this period. The scaled mass index (SMI) was used to determine body condition from body mass and SVL following Peig & Green (40;41). The SMI slope is calculated from the regression of log transformed SVL and log transformed mass.

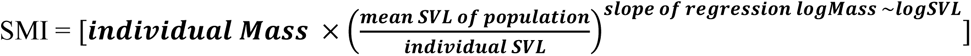

##### CORT assays

After 48 h of acclimation to either 4°C or 8°C, we measured CORT levels in 30 toads using the established water-borne assay protocol by Gabor et al. (42). Briefly, toads were kept in freshly cleaned (EtOH) plastic containers with a volume of 480 mL containing 100 mL of aged and filtered tap water for 1 h. For each sampling batch (i.e., measurement of five toads at the same time), a control sample was run with the sampling water to account for any potential background hormonal traces (43). After taking the control samples the containers were placed in a water bath (Lauda ECO RE 2025 G; LAUDA DR. R. WOBSER GMBH & CO. KG; Lauda-Königshofen, Germany) set to respective acclimation temperature (i.e., 4° and 8°C). During the collection period, nobody was allowed to enter the room to avoid any disturbance of the animals (minimal noise and visual disturbances). After the hour-long sample collection period, we determined the body mass of each toad, which was followed by metabolism measurements.

All water hormone samples were stored at − 25°C. We processed them in a random order within one month. Control samples (N=6) were pooled before processing. Thawed samples were first filtered with Q8 Whatman filter paper to remove suspended particles, and then filtered through C18 solid-phase extraction columns (Oasis Vac Cartridge HLB 3 cc/ 60 mg, 30 µm; Waters, Inc., Switzerland) with a vacuum manifold (Visiprep Vacuum Manifold; Sigma-Aldrich, Germany). The manifold was cleaned before each use with 4 mL of HPLC-grade ethanol and 4 mL of nanopure water. The columns were returned to the −25°C freezer until hormones were eluted with 4 mL of HPLC-grade methanol with a vacuum manifold (Visiprep Vacuum Manifold; Sigma-Aldrich, Germany), which was cleaned with ethanol and nanopure water between batches again. During this process, samples were transferred into 5 mL Eppendorf tubes. Afterwards the methanol was evaporated using a sample concentrator (Stuart sample concentrator, SBHCONC/1; Cole-Parmer, United Kingdom) under a fine N_2_ stream at 45°C using a block heater (Stuart block heater, SBH130D/3; Cole-Parmer, United Kingdom). Dried samples were stored at –25°C until Enzyme-Immunoassay analysis (EIA). The dried samples were re-suspended in a total volume of 500 μL (C. Gabor; personal communication) consisting of 5% ethanol (95% lab grade) and 95% EIA buffer. After re-suspension, samples were frozen at −25°C until vel measurements of hormonal levels via EIA, which took place in May 2022.

The hormonal levels were measured using DetectX Corticosterone ELISA (Enzyme Immunoassay) kits purchased from Arbor Assays (K014-H5, Ann Arbor, MI, USA; assay has a range of 19.53–5,000 pg Corticosterone/mL). Control samples were run on each plate in duplicate. This assay has been previously validated for anurans such as *R. temporaria* (43;44). Samples and kit reagents were brought to room temperature and vortexed before plating. We measured corticosterone concentration in duplicates for all samples on 96-well plates according to the kit’s instructions. The plates were read with a Tecan Spark® Microplate Reader at 450 nm (Tecan, Switzerland). In total, we ran one plate.

Control samples and negative controls did not show CORT levels at detectable ranges. We used MyAssays online tools to calculate the hormonal concentration of samples (https://www.myassays.com/arbor-assays-corticosterone-enzyme-immunoassay-kit-improved-sensitivity.assay). Two positive control samples were run in duplicate on the plate to assess intra-assay variation. Intraplate variation was overall 1.74% (high: 2.68, low: 0.99). The coefficient of variation of duplicates for all samples was 2.72%.

Following Gabor et al. (42), we multiplied CORT release rates (pg × mL^−1^) by the volume of the re-suspension solution (0.5 mL) and standardized values by dividing by the body mass of each individual, resulting in CORT release rates units being pg × g^−1^ × h^−1^.

##### Metabolism and energy expenditure

Oxygen consumption was measured by closed respirometry. Individuals were measured in airtight plastic containers with a volume of 480 mL at their respective acclimation temperature (i.e., 4°C and 8°C). In each container, a planar oxygen sensor spot (SP-PSt3-NAU-D5-YOP, PreSens Precision Sensing GmbH, Regensburg, Germany) was integrated, which was connected to a multichannel oxygen measuring system (Oxy-4 SMA; PreSens Precision Sensing GmbH, Regensburg, Germany) via a fiber optic sensor (Polymer Optical Fiber POF, PreSens Precision Sensing GmbH, Regensburg, Germany). Prior to each trial, the O_2_ fiber optic sensors were calibrated using air-saturated water and a factory-set zero oxygen calibration point at respective acclimation temperature (i.e., 4°C and 8°C) following the calibration protocol of the manufacturer. A temperature probe enabled temperature compensation for dissolved oxygen measurements. The O_2_ concentration was recorded every second and measured as O_2_ % a.s.. Oxygen consumption was measured for 60 min in each animal. Empty (control) chambers were run simultaneously in every trial and values were adjusted accordingly. All trials were conducted between 2000 and 0100 hours to avoid circadian effects. At the end of the oxygen consumption measurements, each animal was placed back in its individual container and measurements of cold tolerance followed.

Oxygen consumption in O_2_ % a.s. was converted to vol % O_2_ and standardized by dividing by the body mass of each individual, resulting in SMR units being mL O_2_ × h^−1^× g^−1^. The analyses were performed in PreSens Oxygen Calculator Software (PreSens Precision Sensing GmbH, Regensburg, Germany).

In order to determine differences in the energy expenditure of the animals under the different acclimation temperatures, DEE in J × (d × g) ^−1^ was calculated from the standardized mean of the metabolic rates per hour and gram of body mass using the caloric equivalent and assuming a respiratory quotient of 0.85 (resulting in 20.37 J × mL O_2_ ^−1^). Finally, we calculated the maximum survival time in d if toads actually relied on the energy storages of the fat bodies to cover their DEE (i.e., energetic maintenance costs) (see 2.3.3).

##### Cold tolerance

The cold tolerance was determined for each toad by using the loss of righting response as a proxy for the lower critical thermal tolerance (CT_min_) according to the procedure of McCann et al. (45). The initial body temperature of the toads was measured by inserting a temperature probe (ThermoPro TP622 IP65 waterproof digital thermometer, ThermoPro DE; precision: ± 0.3 °C) into the animal’s cloaca (i.e., T_cloaca_). The toad was then placed inside a closed plastic container. The container was placed inside an insulated box that was filled with ice. During the cooling, the toad was gently removed from the box every 10 min and placed on its back on a flat surface. If the toad righted itself within 30 s, it was returned to the ice-filled box for further 10 min. After every 10 min the procedures of the measurements were repeated until the toad was no longer able to turn over within 30 s. The T_cloaca_ of the toad at this point was recorded as its CT_min_. This measurement represents a biologically valid and ethically acceptable measure of an animal’s ability to function (45;46). After these procedures, the toads were placed back into their boxes and warmed to their initial acclimation temperature with a heating rate of 0.1°C per minute. All toads recovered quickly.

#### 2.2.3 Statistics

For all statistical tests Cran R (Version 4.1.1, R Development Core Team 2021) for Windows was used. All plots were constructed using ggplot2 (47). Statistical significance was accepted at α < 0.05.

All dependent variables in the models were log_10_-transformed and tested for autocorrelations using Spearman’s rank correlation before the analyses (*cor.test* function in the corTest package; 48). Consequentially, variables were included in statistical analyses when the correlation was significant but below the standard threshold of 0.7 or particularly essential for comparison with previous studies (i.e., SMR/CORT; 49) (Table S3).

Non-parametric Mann-Whitney U-tests were used to compare CORT release rates, SMR, DEE, and CT_min_ between toads acclimated to 4° and 8°C. Paired-samples t-tests were employed to assess the difference in body mass and SMI at collection and at the time of measurements (i.e., after 48 h) of each individual.

### 2.3 Experiment 2: Spring freeze scenario

#### 2.3.1 Animal husbandry, acclimation conditions, and freeze simulation

To explore the physiological responses of *B. bufo* to sudden night freeze during the reproductive period (Table S2), 10 animals were randomly divided into two groups and kept under the same conditions as in experiment 1 for 48 h.

After the 48-h acclimation period, half of the toads (n=5; *freeze group*) were transferred to a portable freezer (DOMETIC Coolfreeze CDF 36, Dometic Germany GmbH, Emsdetten, Germany) and kept at a mean temperature of –2 °C for 6 hours to simulate a night freeze. The other half of the toads remained at 4°C (n=5; *control group*). After 1, 3, and 6 h, we quickly measured T_cloaca_ of all animals by inserting a temperature probe into the animal’s cloaca in their respective acclimation environment (Fig. A1). Toads were handled wearing insulating gloves.

Immediately after freeze exposure, animals from both the freeze group and the control were euthanized by a sharp blow on the head, followed by rapid decapitalization following the protocol of the German animal protection law in an approved laboratory space. Right after euthanizing, the liver was dissected free from surrounding tissue, aliquoted into triplicates, transferred to 1.5 mL Eppendorf tubes filled with RNAlater, snap frozen in liquid nitrogen, and stored at −80°C until transcriptomic analysis.

#### 2.3.2 Blood glucose level

In freshly euthanized toads, blood glucose levels (mg × dL^−1^) were determined in three toads per treatment using a portable kit (CONTOUR®NEXT, Ascensia Diabetes Care Deutschland GmbH, Leverkusen, Germany) from the carotid artery. A Mann-Whitney U-test was used to compare blood glucose levels between toads exposed to freeze and the control group.

#### 2.3.3 Fat body analysis

Fat bodies of four toads were dissected, dry blotted, and weighed to the nearest 0.001 g, followed by drying at 50°C for 24 h. Gross energy content of dissected fat bodies in J was identified by bomb calorimetry (PARR 6100 bomb calorimeter, PARR Instruments Deutschland GmbH, Frankfurt, Germany) with benzoic acid as calibration standard at the University of Hamburg. We lost one sample due to a misfire of the calorimeter. All results were converted from dry matter to fresh weight by correcting the weight loss of the fat body during drying.

#### 2.3.4 Transcriptomics

Total RNA was extracted with a phenol-chloroform precipitation protocol and measured with a NanoDrop 1000 (Peqlab Biotechnologie GmbH, Erlangen, Germany). The RNA samples were then prepared with the Illumina TruSeq stranded mRNA protocol. Afterwards, the samples were quantified with the Quant-iT™ dsDNA BR Assay Kit on a NanoDrop 3300 fluorometer. Equimolar amounts of the samples were pooled. The complete library was run on an Agilent Bioanalyzer prior to being prepared for a NextSeq run, as recommended by Illumina. The library was sequenced on a NextSeq500 using the NextSeq 500/550 HighOutput Kit v2.5 2×75bp cycles sequencing chemistry.

##### Transcriptome assembly & differential expression analysis

The *de novo* assembly of adapter- and quality-filtered single-end reads was carried out using Trinity V.2.8.5 (50;51) and Bowtie (52). Subsequently, the quality and integrity of the assemblies were assessed through the Trinity tool *Trinity.stats*. For the differential expression (DE) analysis, transcript expression levels were quantified using the tool *Salmon* (53), which was integrated into the Trinity pipeline through the execution of the Perl script *align_and_estimate_abundance.pl*. Following this, transcript count matrices were generated, and cross-sample normalization was performed using the *abundance_estimates_to_matrix.pl* script.

Differential expression analysis, conducted at both the gene and isoform levels, was carried out using DESeq2 (54), also integrated into the Trinity pipeline. The p-value threshold was set to p < 0.01, and a minimum count threshold of two was applied. The DESeq output was then employed to create heatmaps for differentially expressed genes and isoforms using the R package *pheatmap* (55). Additionally, volcano plots were generated by plotting the log2 fold change (logFC) against the −log10 (adjusted p-value) using the R package *ggplot2*.

##### Transcriptome annotation

The *de novo* assembly was annotated following the Trinotate pipeline (56) by extracting predicted coding regions using *TransDecoder* (http://transdecoder.github.io) and searching both the entire transcripts and the predicted coding regions separately by BLAST+ (57) against the Uniprot Swiss-Prot database (58), with a stringent expectation value (e-value) of 1 × 10^−5^, retaining only the best BLAST hits. The results were stored in an SQL database, with two distinct tables created for differentially expressed genes and isoforms.

##### Functional classification and gene ontology (GO) enrichment analysis

For functional classification, each BLAST hit was examined, and genes and isoforms with no or contradictory BLAST hits were filtered out. Subsequently, protein functions were identified by referencing the online UniProt database (https://www.uniprot.org/) and conducting a systematic literature review for potential connections between the identified proteins and their roles in freeze response.

The Protein Analysis Through Evolutionary Relationships (PANTHER; 59) Classification System was used to assign Gene Ontology (GO) slim categories, encompassing Biological Processes, Molecular Functions, and Cellular Components, to each differentially expressed protein. Therefore, we extracted the protein IDs from our database and analyzed them in PANTHER to determine their GO slim categories.

Upon obtaining the data from PANTHER, percentage values corresponding to each GO slim category were recorded. To visually present this information, graphics were created in Microsoft Excel, showing the percentage of genes or isoforms assigned to its specific GO slim category. A subsequent step involved plotting the number of genes or isoforms regulated up or down for each biological process category.

## 3. Results

### 3.1 Experiment 1: Warmer winter scenario

There was a significant difference in all physiological parameters between the toads in the 4°C and 8°C acclimation temperature treatment (Table S1; Fig. 1-3). Toads acclimated to 8°C revealed a 3.7-times higher CORT release rate (U=0.00; P<0.001) as well as a 27.1% higher SMR (U=0.0; P<0.001) and used 92.96 J × ^−1^ more energy per day (U=45.00; P=0.004). Mean (±SD) fat body mass and energy content were 0.243 g (±0.08) and 8,652.68 J, respectively. Based on the DEE of the toads at the different acclimation temperatures, mean fat body energy content would be sufficient to cover the energetic maintenance costs 5.26 d longer in toads kept at lower temperatures (i.e., 25.53 d at 4°C; 20.27 d at 8°C; Fig. 1).

**Fig. 1.**
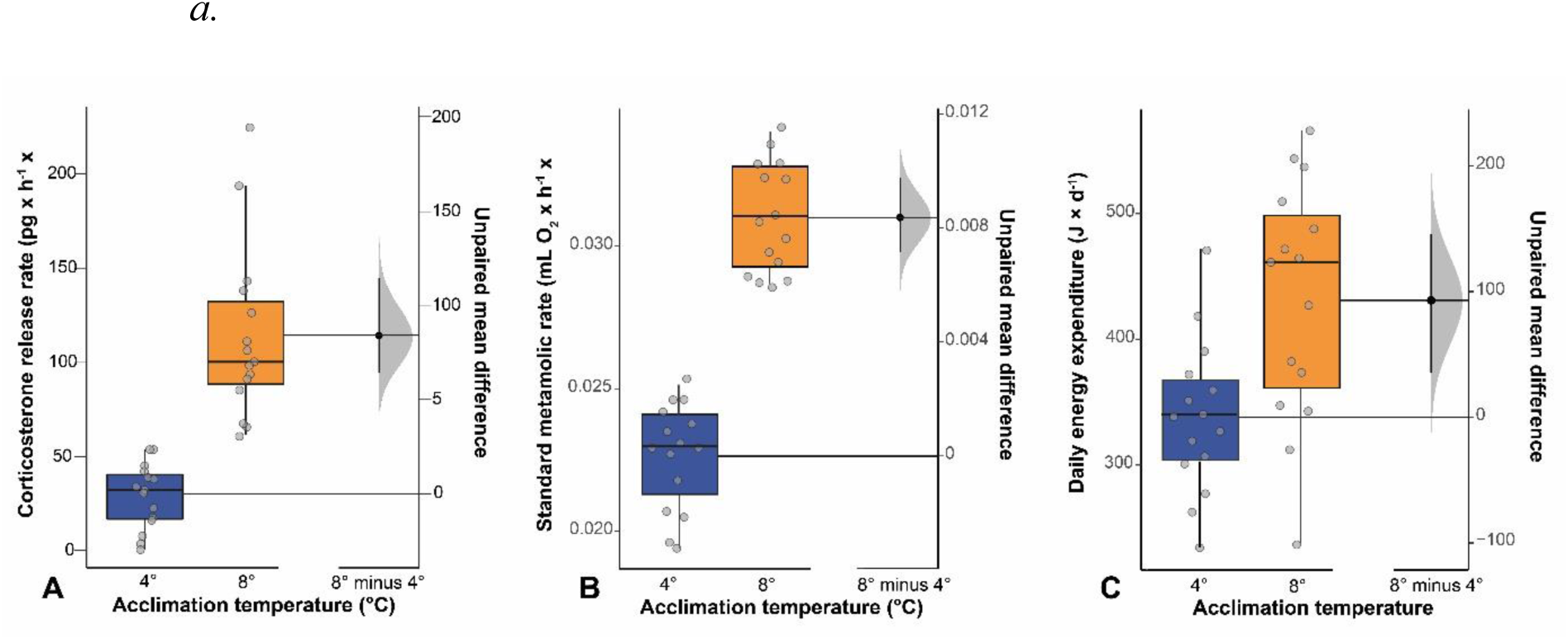
Gardner–Altman estimation plots displaying the mean difference between acclimation temperatures on **A** corticosterone (CORT) release rate (pg × g^−1^× h^−1^), **B** standard metabolic rate (SMR, mL O_2_ × h^−1^× g^−1^), and C daily energy expenditure (J × d^−1^). Boxes and whiskers show 25^th^ to 75^th^ and 10^th^ to 90^th^ percentiles, respectively; black lines indicate the median. Dot=single data points. Blue boxes= toads acclimated to 4°C. Orange box=toads acclimated to 8°C.

There was a significant difference in body mass and body condition before and after the 48-h-long acclimation period (Table S1; Fig. 2) in toads kept at 8°C (body mass: mean= 0.198, SD = 0.02), t(14) = 3.65, P = 0.003; SMI: mean = 0.198, SD = 0.02, t(14) = 3.63, P = 0.003). Body mass and body condition did not change significantly within 48 h in toads kept at 4°C (body mass: mean= −0.004, SD = 0.01), t(14) = −1.322, P = 0.207; SMI: mean = −0.004, SD = 0.01, t(14) = −1.258, P = 0.229). When acclimated to 8°C for 48 h, 73.39% of the toads lost body mass and body condition, whereas 26.61% revealed a higher body mass and body condition.

**Fig. 2.**
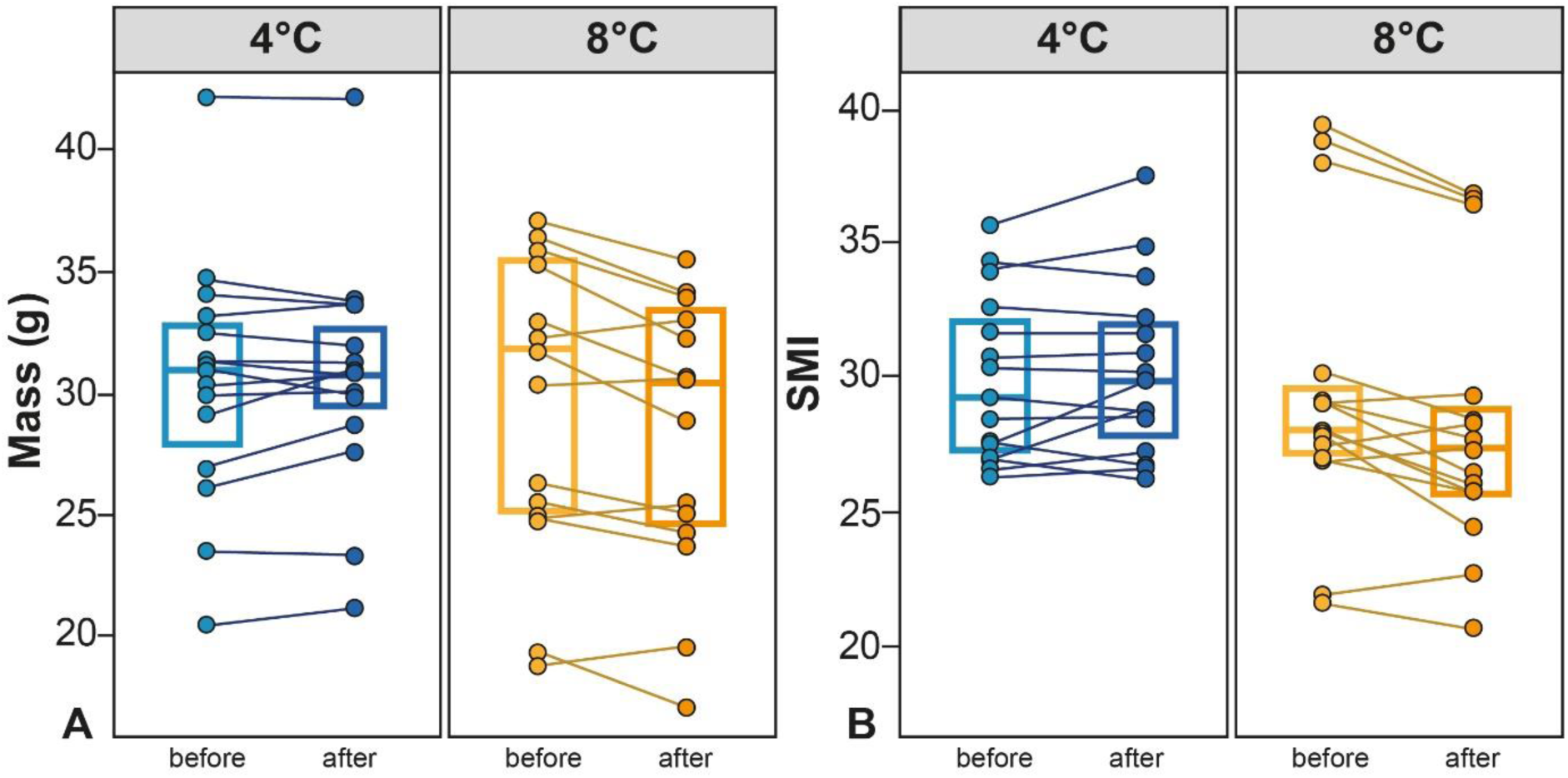
A. Mass (g) and **B** SMI as a measure of body condition of adult *B. bufo* before and after a 48-h acclimation period at 4°C and 8°C. Boxes and whiskers show 25^th^ to 75^th^ and 10^th^ to 90^th^ percentiles, respectively; black lines indicate the median. Dot=single data points. Blue boxes= toads acclimated to 4°C. Orange boxes=toads acclimated to 8°C.

Cold tolerance was significantly lower (i.e., CT_min_ was higher) in toads acclimated to 8 °C (U=2.50; p<0.001; N=30; Fig. 3). Mean (±SD) CT_min_ was −0.167 (±0.13) and 0.34 (±0.14) in toads at 4°C and 8°C, respectively.

**Fig. 3.**
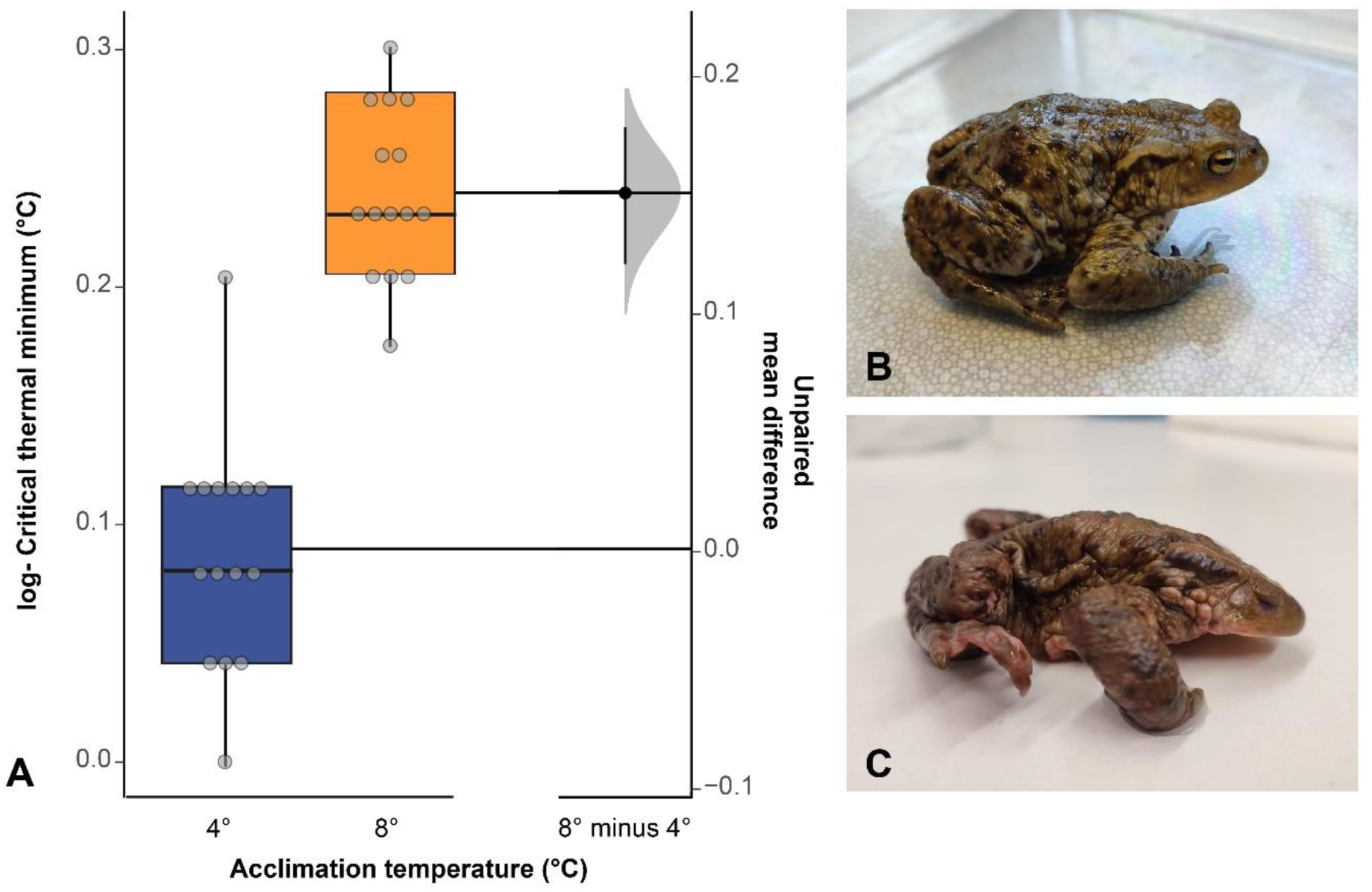
Gardner–Altman estimation plots displaying the mean difference between acclimation temperatures on **A** log-transformed cold tolerance (CT_min_, °C) in adult *B. bufo* (N=30). **B** Start of CT_min_ trial. **C** Toad at CT_min_. Boxes and whiskers show 25^th^ to 75^th^ and 10^th^ to 90^th^ percentiles, respectively; black lines indicate the median. Dot=single data points. Blue box= freeze group. Orange box=control group.

### 3.2 Experiment 2: Spring freeze scenario

#### 3.2.1 *De novo* assembly and annotation

The *de novo* assembly was conducted using RNA-seq data obtained from both control (n = 5) and frozen toad (n = 5) samples. cDNA libraries were constructed from the liver tissue of *B. bufo*, resulting in a comprehensive assembly comprising 231,696 transcripts (see Table S4 for full assembly statistics). Employing DESeq2 for differential gene expression analysis revealed significant variations in both gene and isoform expression patterns between the freeze and control group (Fig. 4, S2-5). A total of 54 differentially expressed transcripts were identified at the gene level, while at the isoform level, 277 transcripts showed distinct expression profiles (Fig. 4, S4). Subsequent annotation using Trinotate, coupled with a stringent filtering process for BLAST results (excluding transcripts lacking BLAST hits and those with contradictory outcomes), allowed us to assign names and functions to six proteins at the gene level and 173 proteins at the isoform level.

**Fig. 4.**
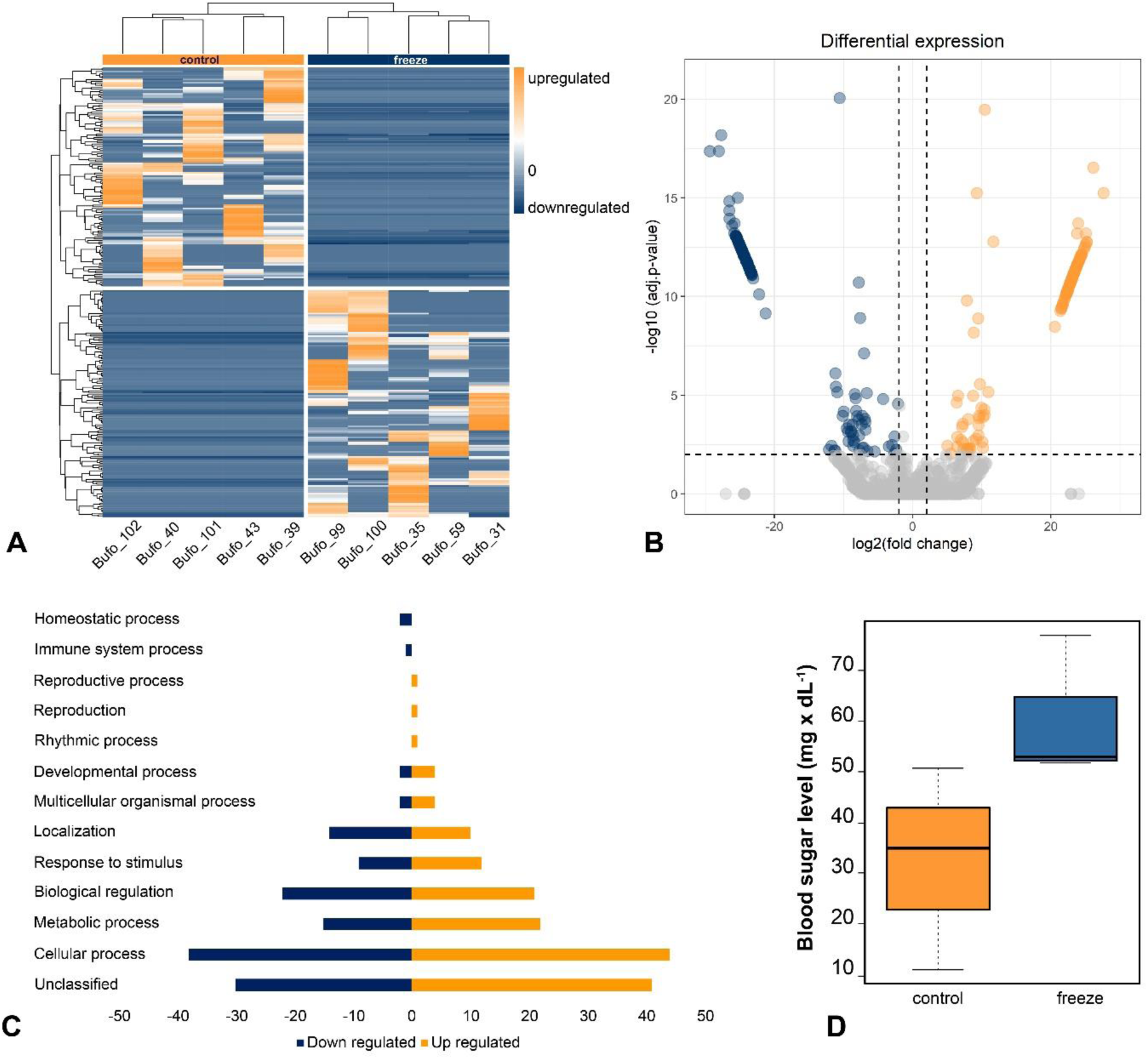
A. Heatmap with hierarchical clustering of genes (rows) showing relative expression of all significant differentially expressed gene isoforms (DEGs) (P<0.01) in individual control group (N=5, left column) and individual freeze group (N=5, right column) livers. Orange indicates upregulation, blue indicates downregulation. **B** Volcano plot showing DEGs in transcriptomes of control group and freeze group livers. The logFC in counts for each transcript (magnitude of differential expression) are plotted on the x-axis against the statistically significant −log10 (P_adj_ <0.01) values on the y-axis. Dashed lines correspond to P_adj_ and logFC cut-off values set at P<0.01 and 2.0, respectively. Grey dots indicate DEGs (P_adj_ <0.01); orange and blue dots indicate DEGs (P_adj_ <0.01) above or below the set logFC threshold of +2.0 or −2.0, respectively. **C** Biological process categories of isoforms differentially expressed in *Bufo bufo* compared to the control group. Data show the number of/percentage of differently expressed gene isoforms that were up- and downregulated in frozen toads. **D** Effect of freeze exposure on blood sugar level (mg × dL^−1^). Boxes and whiskers show 25^th^ to 75^th^ and 10^th^ to 90^th^ percentiles, respectively; black lines indicate the median. Dot=single data points. Blue box= freeze group (n=3). Orange box=control group (n=3).

#### 3.2.2 Differential gene expression

As only six out of 54 differentially expressed transcripts on gene level between frozen and control toads were identified, the analysis herein will focus on comparisons on isoform level. We present the analysis of the gene level in the electronical supplementary material (Fig. A2–4; Table S5). Isoform-level analysis revealed 277 transcripts that were differently expressed (Fig. 4A-C; Fig. A5; Table S6). Using isoform-level analysis, exposure to freeze resulted in downregulation of 136 (i.e., 84 annotated) transcripts and upregulation of 141 (i.e., 89 annotated) transcripts relative to control animals (Fig. 4A-C).

#### 3.2.3 Functional classification of differentially regulated genes

The biological process-oriented functional classification identified 26.9% of isoforms associated with cellular processes (Fig. A5). Furthermore, 13.8% of isoforms were categorized as related to metabolic processes and 14.2% to biological regulation. The most common molecular functions of isoforms were *catalytic* (26.2%) and *binding* (27.4%). Most of isoforms were associated with the cellular anatomical entity (53.9%).

Exposure to freeze resulted in differential expression of isoforms related to several GO slim Biological Process categories including amino-acid biosynthesis, carbohydrate metabolism and particularly gluconeogenesis as well as lipid metabolism, and stress (Fig. 4A-C, Table S6).

Differently expressed gene isoforms associated with cellular (up: 45.83%; down: 39.58%) as well as metabolic processes (up: 46.81%; down: 31.91%) and responses to stimuli (up: 44.44%; down: 22.22%) in toads exposed to freeze were mostly upregulated relative to control animals. Gene isoforms related to regulative biological regulation were shared equally up- and downregulated (up: 42%; down: 44%).

Among the upregulated gene isoforms in toads exposed to freeze were terms associated with carbohydrate and/or lipid metabolism were cryptochrome-2 (CRY2) and insulin-like growth factor-binding protein 4 (IGFBP4). Similarly, a significant increase in the expression of aspartate aminotransferase (GOT1), involved in amino-acid biosynthesis, urea cycle, and gluconeogenesis, was shown in frozen toads compared to the control animals. Another notable upregulated isoform was adenylate cyclase type 9 (ADCY9), known to contribute to signaling cascades activated by corticotropin-releasing factor, corticosteroids and thus, involved in the neuroendocrine response to stress.

In contrast, several gene isoforms associated with carbohydrate and/or lipid metabolism were downregulated during freeze exposure, namely fructose-1,6-bisphosphatase 1 (FBP1), C2 domain-containing protein 5 (C2CD5), glutamine-fructose-6-phosphate aminotransferase 1 (GFPT1), beta-2-glycoprotein 1 (APOH), and cytoplasmic polyadenylation element-binding protein 4 (CPEB4). Furthermore, N-acetylglutamate synthase (NAGS), involved amino-acid biosynthesis and urea cycle, was downregulated.

#### 3.2.3 Blood glucose level

Mean blood glucose level was 83.51% higher in frozen toads compared to the control group (U=0.05; P=0.061; N=6; Fig. 4D).

## 4. Discussion

### 4.1 The Toll of Warmer Winter Days: Increased Metabolic Demands and Reduced Cold Tolerance

In ectothermic animals such as amphibians, metabolic rate and thus, energy demands increase with ambient temperature (60). This may have implications for animal health and survival, particularly when resources are limited, such as during winter. Here, we observed that toads exposed to a 4°C warmer temperature had increased metabolic rates and higher daily energy demands, leading to a faster depletion of energy budgets over time compared to toads kept at standard winter conditions (i.e., 4°C). Across vertebrates, increased energy demands resulting from homeostatic challenges such as adverse environmental conditions are met through the release of glucocorticoid hormones (i.e., CORT in amphibians; 61) via the activation of the neuroendocrine stress axis/hypothalamic–pituitary–adrenal/interrenal (HPA/I) axis (62;63). Glucocorticoid hormones play a pivotal role in the metabolism of most energy reserves, regulating glucose, fat, and protein metabolism (63;64). In response to increases in metabolic rate, a main role of these hormones is to increase circulating glucose levels at a rate that corresponds to metabolic demands (62;65). Thus, higher metabolic activity is suggested to be concomitant with increases in CORT levels (66). Indeed, in toads kept at a 4°C warmer temperature, we observed higher CORT release rates, suggesting that the temperature-induced higher energy demands observed therein was met through the mobilization of energy reserves mediated by an increased secretion of the glucocorticoid hormone. Furthermore, energy storages decreased withing the 48 h of exposure to higher temperatures, indicating that these reserves were mobilized to meet the increased energy demands. Consequently, warmer winter days will result in changes in overall energy expenditure during overwintering, leading to accelerated depletion of energy reserves for all ectotherms that rely on internal energy reserves, not just amphibians (22;23;67). Here, for instance, we found that the fat body, as the major internal energy reserves in overwintering amphibians, would be depleted 5.26 d earlier in toads kept at higher temperatures. Despite the significant reduction in survival probabilities during overwintering and at the beginning of the breeding season upon leaving the overwintering shelter because of starvation, depleted energy reserves may further reduce immune system functioning and fecundity. This, coupled with elevated CORT levels, might increase their vulnerability to other stressors (25;68;69;70).

In our study, cold tolerance was remarkably lower in toads kept at a 4°C warmer temperature, presumably linked to the change of energy expenditure experienced by these toads as well as the onset of physiological acclimation to higher temperatures (4;71;72). Cold tolerance and energetics are closely related, as a decrease in energy reserves reduces the ability to respond effectively to cold stress, given that those energy reserves are required to cover the energy expended during cold stress exposure (6). In freeze-tolerant species, carbohydrate energy reserves are essential for survival of freezing itself in their function as cryoprotectants (34;35). Our results are in line with findings for other ectothermic species with a life-stage specific tolerance to winter conditions. For example, Sobek-Swant et al. (73) found that warmer winter temperatures resulted in the loss of winter acclimatization in the emerald ash borer (*Agrilus planipennis*), thereby increasing their vulnerability to cold spells. Furthermore, in the freeze-tolerant wood frog (*Lithobates sylvaticus*), repeated freeze–thaw due to increased winter temperature variability reduced energy reserves and thereby cold tolerance (74). Therefore, changing thermal winter conditions are likely to result in a mismatch between the occurrence of extreme thermal events such as sudden freeze and the physiological mechanisms to tolerate them (4;75;74). As a result, ectotherms including amphibians might be more susceptible to cold stress.

### 4.2 Toads on Ice: Limited Physiological Responses to Sudden Freeze

Amphibians inhabiting seasonal cold climates have developed a variety of winter survival strategies such as overwintering underwater or underground. However, some species that overwinter at or near the soil surface, have evolved a freeze tolerance as an adaptive cold hardiness strategy that allows surviving harsh thermal conditions (rev. in 35). When exposed to freezing temperatures, these amphibians can synthesize and accumulate high concentrations of cryoprotectants such as glucose, glycerol, or urea in their tissues (rev. in 35). For example, the synthesis of glucose in *L. sylvaticus* is triggered by ice crystallization on the skin followed by a homeostatic stress response that stimulates glycogenolysis in the liver and its export to all other organs, resulting in a hyperglycemic state (34). To that end, carbohydrate energy reserves (i.e., glycogen) are mobilized and enzymes associated with liver glycogenolysis are upregulated (76). In our study, the blood glucose level was indeed 83% higher in frozen toads. Even if this difference was not significant, probably due to a low statistical power resulting from a small sample size, this result suggests the mobilization of carbohydrate reserves. In addition, we found evidence of transcriptional regulation of glucose biosynthesis pathways. However, contrary to our expectations, that response did not entail enhanced expression of gene isoforms directly promoting glucose synthesis via hepatic glycogenolysis (76). Those enzymes, including glycogen phosphorylase (GP) and glucose-6-phosphatase (G6PC), did not show differential expression in frozen toads, and glucose transporters required for the export of glucose from the liver such as GLUT2 (35;76), were not responsive to freezing conditions in this study. Furthermore, in the pathway for glucose biosynthesis through gluconeogenesis, fructose-1,6-bisphosphatase 1 (FBP1), as one of the key enzymes, was diminished in frozen toads (35;76). Nonetheless, we found that other notable gene isoforms associated with glucose metabolism and homeostasis, mostly in response to starvation, were upregulated in frozen toads. For example, aspartate aminotransferase (GOT1) regulates the uptake of glutamine by the liver, thereby fueling the synthesis of glucose through gluconeogenesis (77). Moreover, glutamine-fructose-6-phosphate aminotransferase 1 (GFPT1) was downregulated in response to freezing. Downregulation of GFPT1 might contribute to glucose availability for accumulation as it prevents the flux of glucose from the anaerobic energy production pathway to the hexosamine pathway (78). These findings suggest a response to starvation/fasting or reduced glucose availability (77;79) to guarantee the availability of glucose for cryoprotective purposes. This might be further supported by adenylate cyclase type 9 (ADCY9), an enzyme involved in signaling cascades of the neuroendocrine stress axis and thus, contributing to the role of CORT in the regulation of circulating glucose levels to meet energetic demands (62;65). Therefore, our results indicate that exposure to sudden freeze led to physiological responses to maintain homeostasis as well as an increase of cold hardiness by the mobilization of the cryoprotectant glucose. In contrast, we could not find any evidence for the synthesis of glycerol or urea as additional cryoprotectants as, for example, N-acetylglutamate synthase (NAGS), known to be involved in the urea cycle, was downregulated. Measurements of urea could be included by future studies (80). Consequently, *B. bufo* might be capable of at least a short-term freeze tolerance which might provide a means in fitness if toads encounter a sudden night freeze, for instance, on their way to their breeding sites after leaving the overwintering shelter. Nonetheless, the mobilization of glucose for cryoprotective purposes requires energy reserves (i.e., carbohydrates for glycogenolysis and proteins for gluconeogenesis). If these energy reserves are already drained due to the catabolic effect of increased mean winter temperatures with climate change (6), cryoprotective accumulation of glucose might be constrained with consequences for animal survival in false springs events. Future research is needed to better understand the interplay of elevated mean temperatures and an increased frequency of extreme cold events on amphibian physiological resilience. This research is crucial to unraveling the potential double jeopardy of winter climate change, which may have been underestimated in its role in the decline of temperate amphibian populations.

## 5. Conclusion

Energy is the only currency that counts in and for life, and climate change has the potential to disrupt energy balances throughout the annual cycle in several ways such as through increased energetic demands and fewer available resources. Consequently, wildlife might become more susceptible to environmental stress, particularly when coping mechanisms are limited. Our study points for the first time to the complex physiological challenges that amphibians face as a result of changing thermal conditions due to winter climate change. Overall, our results highlight that warmer winters and repeated freeze-thaw cycles impose physiological costs that could reduce energy reserves and might thereby affect amphibian health and survival. Moreover, we demonstrated that warmer winter temperatures reduced cold tolerance, suggesting that both the increases in mean temperatures as well as a period of warm winter days will make amphibians more sensitive to extreme thermal events, which are increasing in frequency as a consequence of climate change. We hope this study will encourage future research further exploring the complex effects of (winter) climate change on wildlife energy budgets and coping mechanisms in order to improve our ability to predict species’ vulnerability to global warming.

## 6. Data Accessibility Statement

All raw data generated and analyzed in this study as well as all the scripts employed in the transcriptomic analysis are available at https://figshare.com/s/8a5b702eacc2aa85cfcc (DOI: 10.6084/m9.figshare.24891765, to be published after acceptance). A preprint has been uploaded to biorvix (DOI: https://doi.org/10.1101/2024.01.03.574103; 81).

## 1. Appendix

**Fig. A1.**
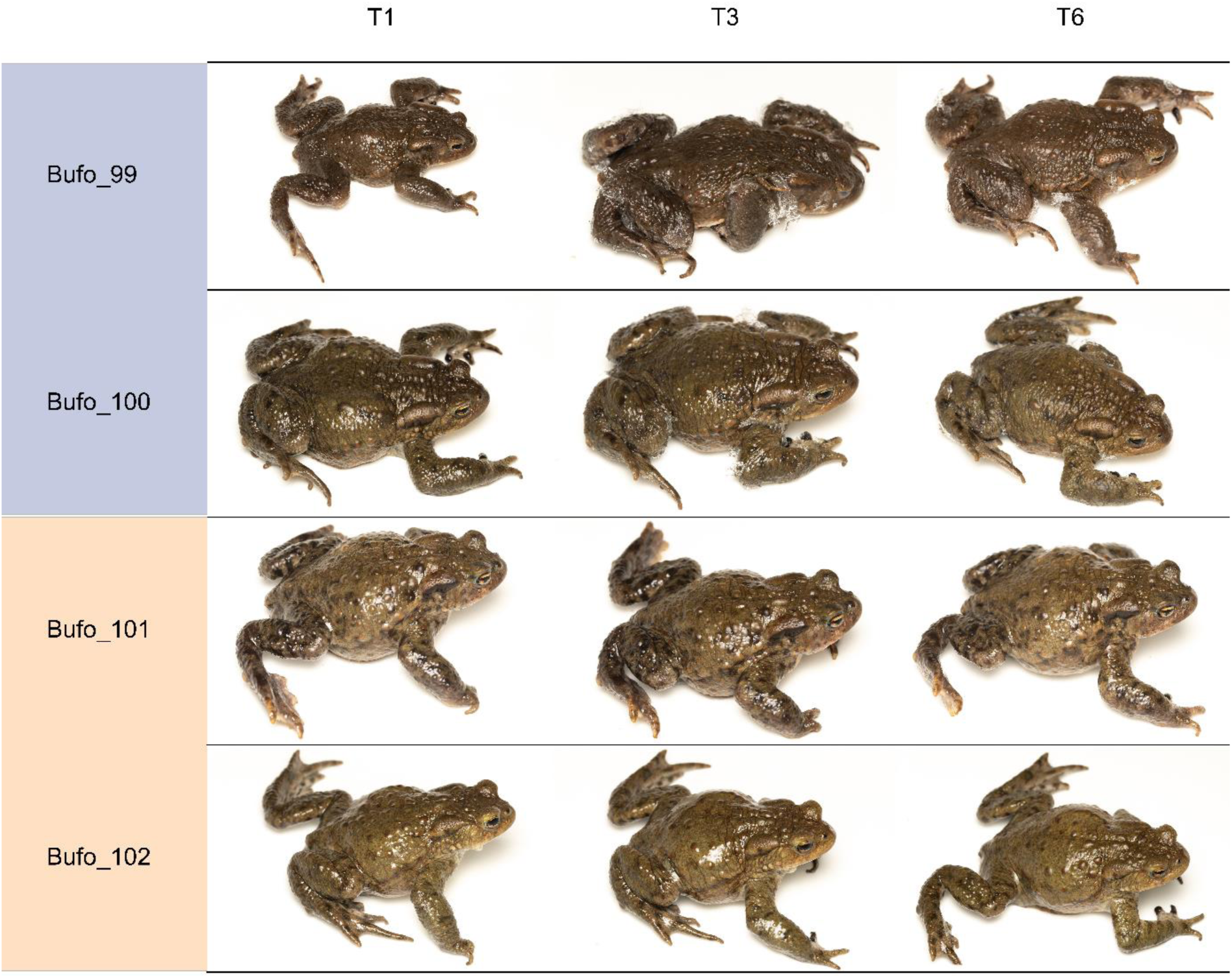
Graphical overview of toads from the control group (orange) and the freeze group (blue) during the 6-hour long freeze simulation. T1= 1 hour after experiment start. T3= 3 hours after experiment start. T6= 6 hours after experiment start/end of the experiment.

**Fig. A2.**
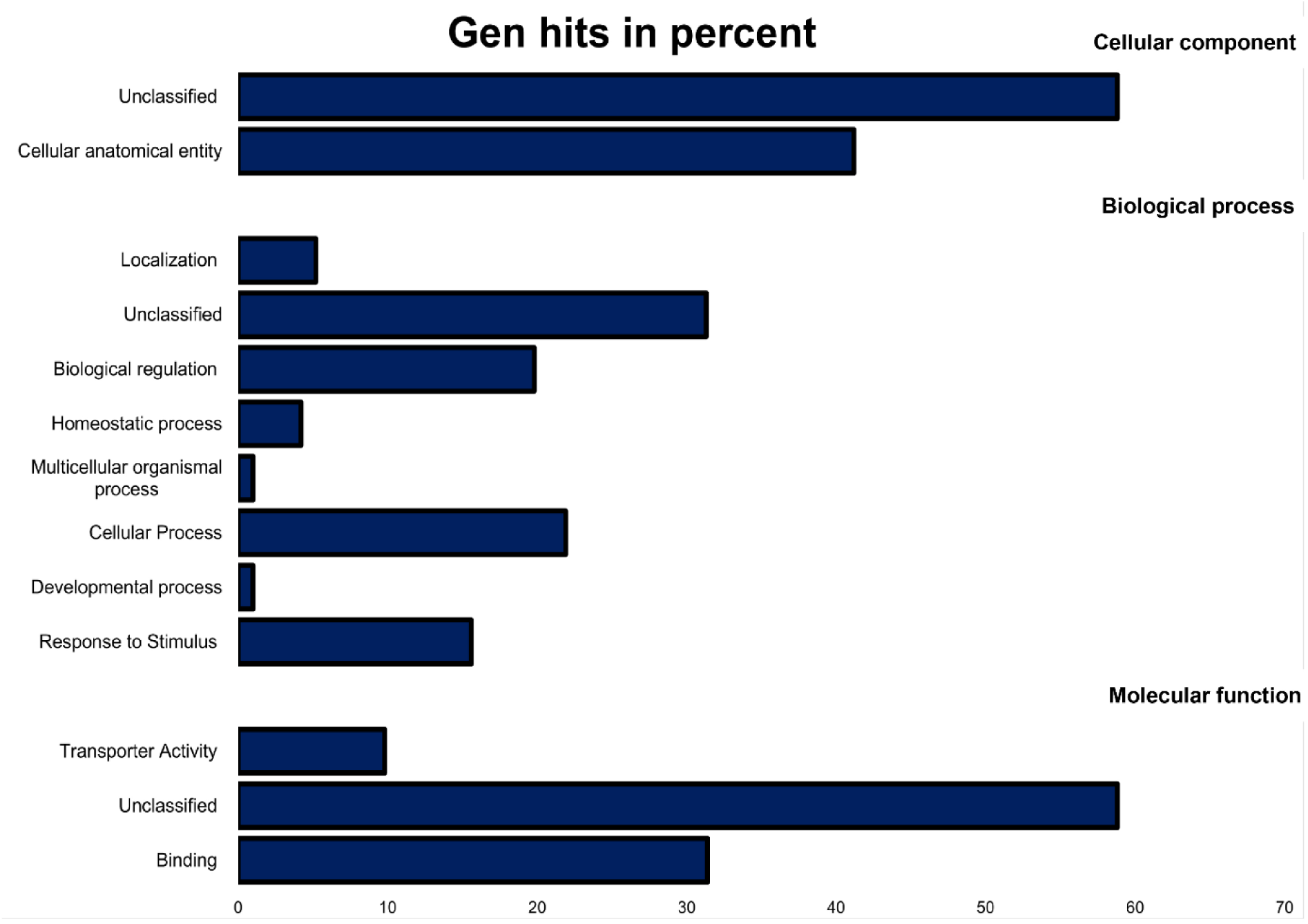
Gene ontology terms describing the differential expressed genes that were up- and downregulated in frozen toads in *Bufo bufo*. Data shown are percentages of total genes annotated. Most genes were associated with the cellular anatomical entity (41.2%). The most common molecular functions of genes were transporter activity (9.8%) and binding (31.4%). The biological process-oriented functional classification of the annotated transcriptome identified 21.9% of genes associated with cellular processes. Furthermore, 15.6% of genes were categorized as related to stimulus responses and 19.8% to biological regulation.

**Fig. A3.**
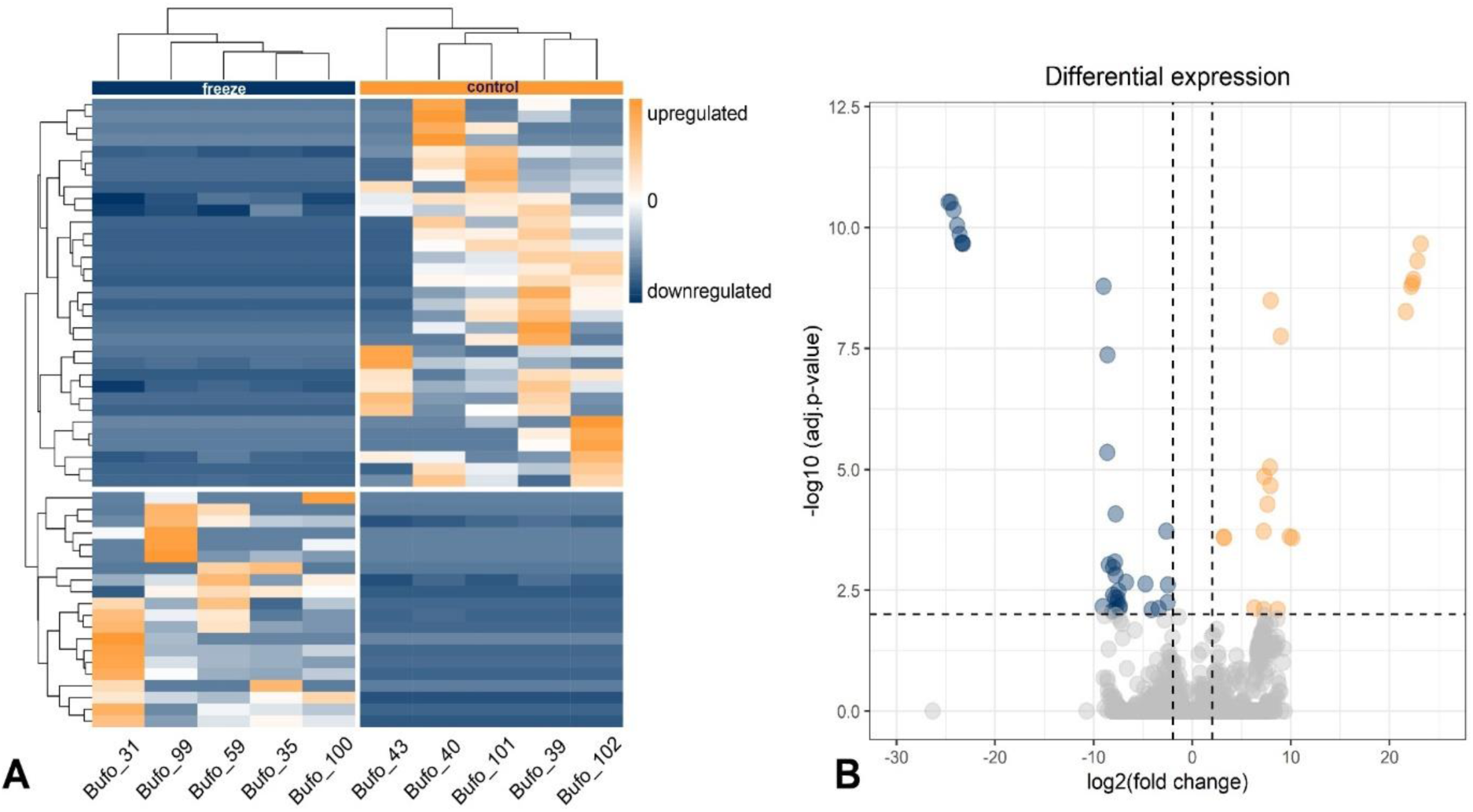
**A**. Heatmap with hierarchical clustering of genes (rows) showing relative expression of all significant differentially expressed genes (DEGs) (P<0.01) in individual control group (N=5, left column) and individual freeze group (N=5, right column) livers. Orange indicates upregulation, blue indicates downregulation. **B**. Volcano plot showing DEGs in transcriptomes of control group and freeze group livers. The logFC in counts for each transcript (magnitude of differential expression) are plotted on the x-axis against the statistically significant −log10 (P_adj_ <0.01) values on the y-axis. Dashed lines correspond to Padj and logFC cut-off values set at P<0.01 and 2.0, respectively. Grey dots indicate DEGs (Padj <0.01); orange and blue dots indicate DEGs (P_adj_ <0.05) above or below the set logFC threshold of +2.0 or −2.0, respectively. Gene-level analysis revealed 54 transcripts that were differently expressed between frozen and control toads. Exposure to freeze resulted in downregulation of 32 (i.e., 2 annotated) transcripts and upregulation of 20 (i.e., 3 annotated) transcripts relative to control animals.

**Fig. A4.**
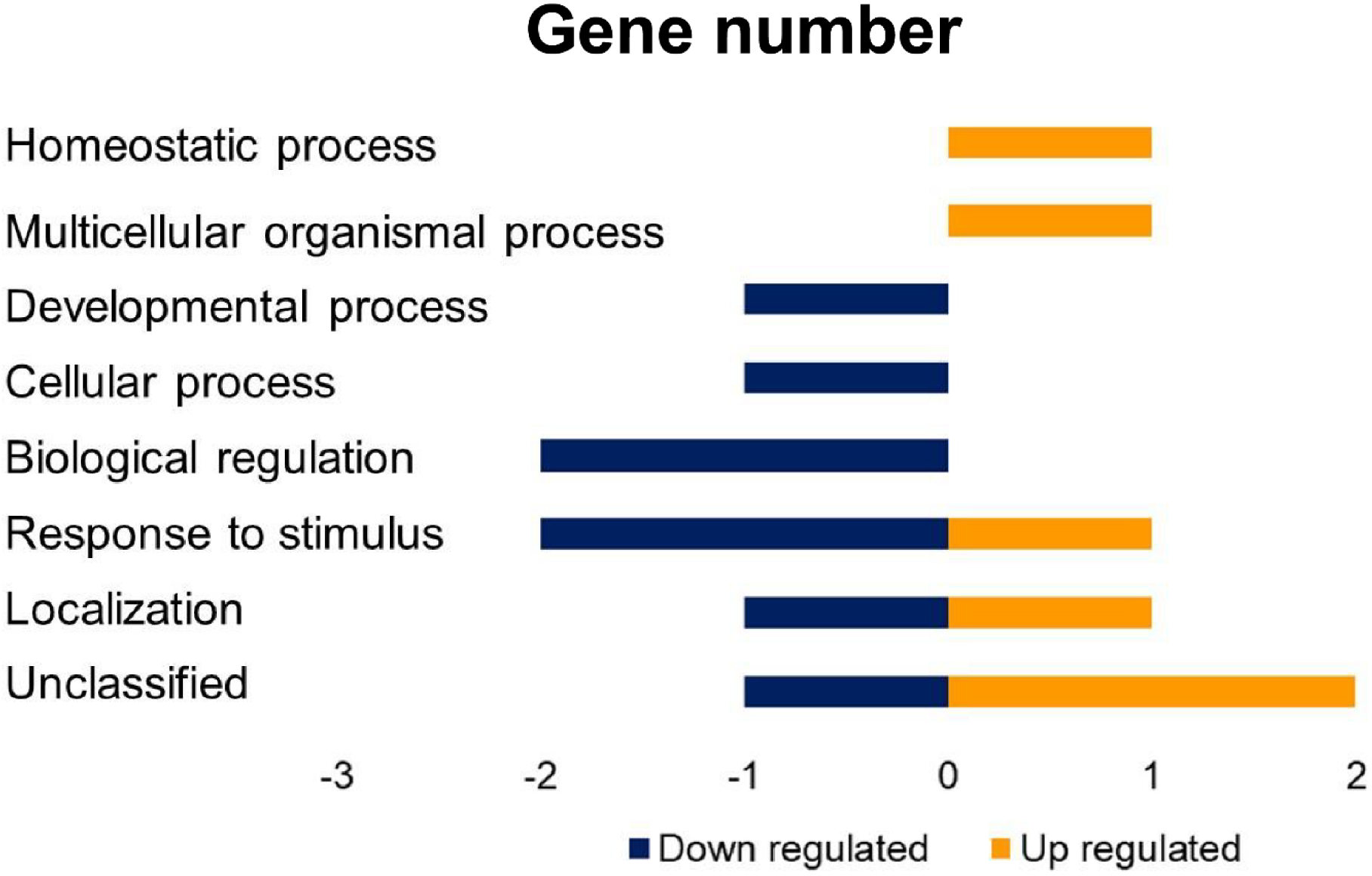
Biological process categories of genes differentially expressed in *Bufo bufo* compared to the control group. Data show the number of differently expressed genes that were up- and downregulated in frozen toads. Gene-level analysis revealed 54 transcripts that were differently expressed between frozen and control toads. Exposure to freeze resulted in downregulation of 32 (i.e., 2 annotated) transcripts and upregulation of 20 (i.e., 3 annotated) transcripts relative to control animals.

**Fig. A5.**
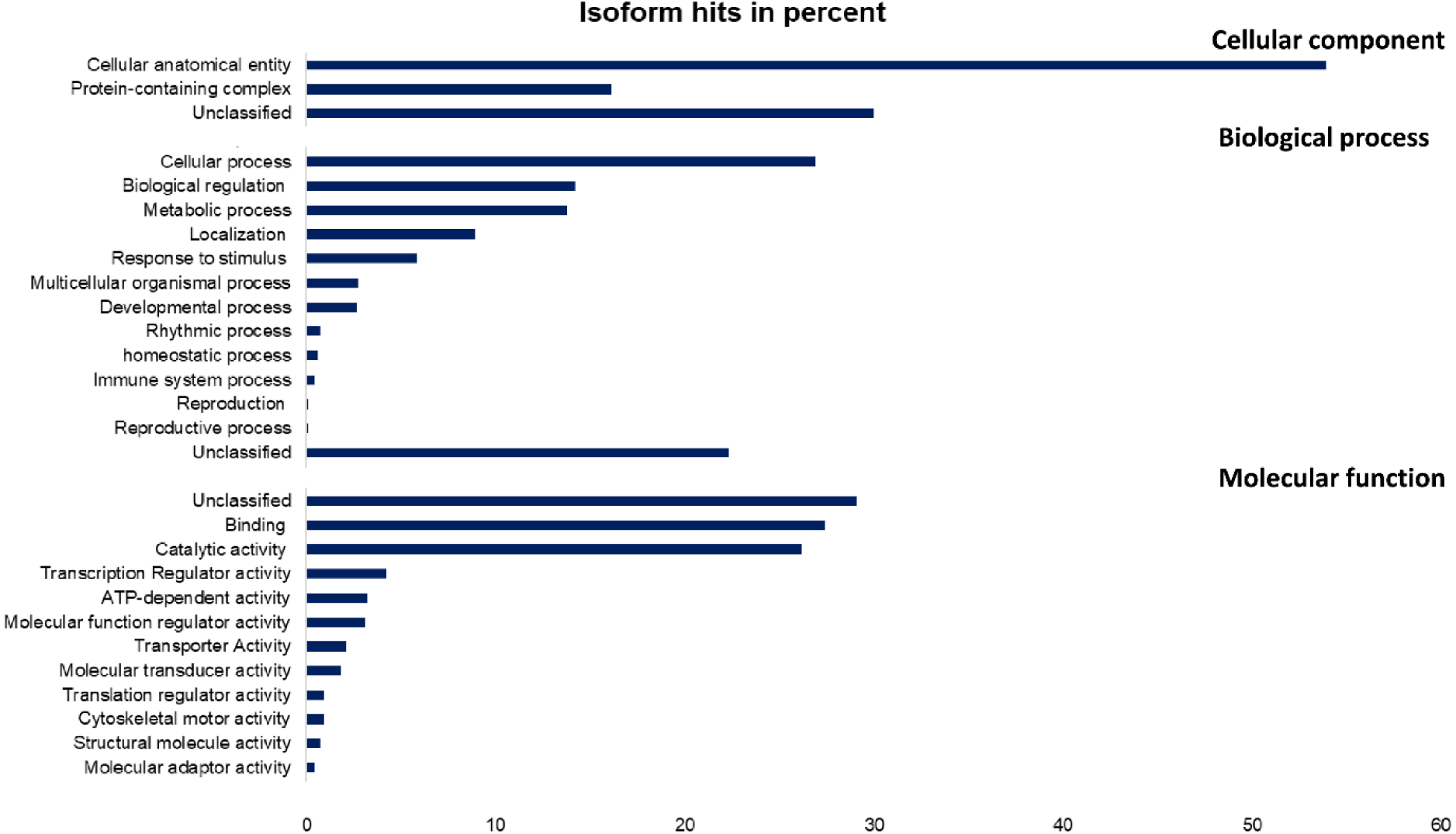
Gene ontology terms describing the differential expressed isoforms that were up- and downregulated in frozen toads in *Bufo bufo*. Data shown are percentages of total isoforms annotated. Most isoforms were associated with the cellular anatomical entity (53.9%). The most common molecular functions of isoforms were catalytic activity (26.2%) and binding (27.4%). The biological process-oriented functional classification identified 26.9% of isoforms associated with cellular processes. Furthermore, 13.8% of isoforms were categorized as related to metabolic processes and 14.2% to biological regulation.

## 9. Acknowledgements

We thank Miguel Vences, Ben Oetken, Sven Gippner, Fabian Bartels, Paula C. Eterovick, and Janina Rudolph for field assistance, Gabriele Keunecke her help in the laboratory in Braunschweig, and Maileen Weidner as well as Sina Remmers for their help in the laboratory in Hamburg. We are grateful to Jelena Mausbach for providing useful methodological information and advice on the water-borne CORT extractions and assays during the establishment of the protocol at the Technische Universität Braunschweig, and Caitlin Gabor for providing helpful information on the CORT assays. We thank Karsten Hiller for providing the possibility for carrying out the CORT assays in the laboratory of the BRICS (Braunschweig Integrated Centre of Systems Biology) and Antonia Henne for her help in the laboratory at the BRICS. We thank Hans-Peter Ruthsatz for providing the blood glucose meter. We are grateful to Julia Nowack, Jérémy Terrien, and Sylvain Giroud for inviting KR to the Centenary Conference of the Society of Experimental Biology 2023 in Edinburgh to present this work in the session AC1-*Thermoregulatory and metabolic adaptations in a changing world*. The Society for Experimental Biology funded the travel expenses of KR.

## 10. Author contributions

**RS:** Conceptualization (equal); Supervision (equal); Methodology (equal); Data curation (equal); Formal analysis (lead); Investigation (equal); Writing – original draft (lead); Writing – review and editing (equal). **CZ**: Investigation (equal); Data curation (supporting); Writing – review and editing (supporting). **NS**: Investigation (equal); Data curation (supporting); Writing – review and editing (supporting). **JSP**: Formal analysis (equal); Writing – review and editing (equal). **SK**: Methodology (equal); Writing – review and editing (equal). **KHD**: Resources (supporting); Writing – review and editing (supporting). **KR**: Conceptualization (equal); Supervision (equal); Methodology (equal); Data curation (equal); Formal analysis (lead); Investigation (equal); Writing – original draft (lead); Writing – review and editing (equal). All authors gave final approval for publication and agreed to be held accountable for the work performed therein.

## 11. Conflict of Interest

The authors declare have no competing interests.

## 12. Statement of Ethics

The authors have no ethical conflicts to disclose. The experiments were conducted under permission from the *Niedersächsisches Landesamt für Verbraucherschutz und Lebensmittelsicherheit*, Germany (Gz. 33.19-42502-04-21/3705). Fieldwork in Lower Saxony was carried out with permits of Stadt Braunschweig (Stadt Braunschweig - Fachbereich Umwelt und Naturschutz, Richard-Wagner-Straße 1, 38106 Braunschweig; Gz. 68.11-11.8-3.3).

## 13. Funding

JSP was supported by the European Union’s Horizon 2020 program under the Marie Skłodowska-Curie grant (agreement ID: 101028000). RS was funded by the DFG priority program SPP 1991 Taxon-Omics (grant number VE247/20-1).

## Notes

### Competing Interest Statement

The authors have declared no competing interest.

### Summary of Updates

Title has been changed. Clarification of some methodological details.

## References

1. Sunday J, Bennett JM, Calosi P, Clusella-Trullas S, Gravel S, Hargreaves AL, Leiva FP, Verberk WCEP, Olalla-Tárraga MÁ, Morales-Castilla I. 2019 Thermal tolerance patterns across latitude and elevation. Philos. Trans. Royal Soc. B, 374, 20190036. (doi: 10.1098/rstb.2019.0036)

2. Comte L, Olden JD. 2017 Climatic vulnerability of the world’s freshwater and marine fishes. *Nat*. Clim. Change, 7, 718–722. (doi: 10.1038/nclimate3382)

3. Morley SA, Peck LS, Sunday JM, Heiser S, Bates AE. 2019 Physiological acclimation and persistence of ectothermic species under extreme heat events. Global Ecol. Biogeogr., 28, 1018–1037. (doi: 10.1111/geb.12911)

4. Williams CM, Henry HA, Sinclair BJ. 2015 Cold truths: how winter drives responses of terrestrial organisms to climate change. Biol. Rev., 90, 214–235. (doi: 10.1111/brv.12105)

5. Lee H, Calvin K, Dasgupta D, Krinner G, Mukherji A, Thorne P, Trisos C, Romero J, Aldunce P, Zommers Z, et al. 2023 Climate change 2023: synthesis report. Contribution of working groups I, II and III to the sixth assessment report of the intergovernmental panel on climate change. (doi: 10.59327/IPCC/AR6-9789291691647)

6. Marshall KE, Gotthard K, Williams CM. 2020 Evolutionary impacts of winter climate change on insects. Curr. Opin. Insect Sci., 41, 54–62. (doi: 10.1016/j.cois.2020.06.003)

7. Cohen J, Screen JA, Furtado JC, Barlow M, Whittleston D, Coumou D, Francis J, Dethloff K, Entkhabi D, Jones J, et al. 2014 Recent Arctic amplification and extreme mid-latitude weather. Nat. Geosc., 7, 627–637. (doi: 10.1038/ngeo2234)

8. Wallace JM, Held IM, Thompson DW, Trenberth KE, Walsh JE. 2014 Global warming and winter weather. Sci., 343, 729–730. (doi: 10.1126/science.343.6172.729)

9. Stendel M, Francis J, White R, Williams PD, Woollings T. 2021 The jet stream and climate change. Climate Change, 327–357. (doi: 10.1016/B978-0-12-821575-3.00015-3)

10. Walther GR, Post E, Convey P, Menzel A, Parmesan C, Beebee TJ, Fromentin JM, Hoegh-Guldberg O, Bairlein F. 2002 Ecological responses to recent climate change. Nature, 416, 389–395. (doi: 10.1038/416389a)

11. Parmesan C, Yohe G. 2003 A globally coherent fingerprint of climate change impacts across natural systems. Nature, 421, 37–42. (doi: 10.1038/nature01286)

12. Visser ME, Both C. 2005 Shifts in phenology due to global climate change: the need for a yardstick. Proc. Royal Soc. B: Biol. Sci., 272, 2561–2569. (doi: 10.1098/rspb.2005.3356)

13. Cohen JM, Lajeunesse MJ, Rohr JR. 2018 A global synthesis of animal phenological responses to climate change. *Nat*. Clim. Change, 8, 224–228. (doi: 10.1038/s41558-018-0067-3)

14. Cleland EE, Chuine I, Menzel A, Mooney HA, Schwartz MD. 2007 Shifting plant phenology in response to global change. Trends Ecol. Evol., 22, 357–365 (doi: 10.1016/j.tree.2007.04.003)

15. Gordo O. 2007 Why are bird migration dates shifting? A review of weather and climate effects on avian migratory phenology. Clim. Res., 35, 37–58. (doi: 10.3354/cr00713)

16. Forrest JR. 2016 Complex responses of insect phenology to climate change. Curr. Opin. Insect Sci., 17, 49–54. (doi: 10.1016/j.cois.2016.07.002)

17. Wells CP, Barbier R, Nelson S, Kanaziz R, Aubry LM. 2022 Life history consequences of climate change in hibernating mammals: a review. Ecography, 2022, e06056. (doi: 10.1111/ecog.06056)

18. Green DM. 2017 Amphibian breeding phenology trends under climate change: predicting the past to forecast the future. Global Change Biol., 23, 646–656. (doi: 10.1111/gcb.13390)

19. Parmesan C. 2007 Influences of species, latitudes and methodologies on estimates of phenological response to global warming. Global Change Biol., 13, 1860–1872. (doi: 10.1111/j.1365-2486.2007.01404.x)

20. Li Y, Cohen JM, Rohr JR. 2013 Review and synthesis of the effects of climate change on amphibians. Integr. Zool., 8, 145–161. (doi: 10.1111/1749-4877.12001)

21. While GM, Uller T. 2014 Quo vadis amphibia? Global warming and breeding phenology in frogs, toads and salamanders. Ecography, 37, 921–929. (doi: 10.1111/ecog.00521)

22. Reading CJ. 2007 Linking global warming to amphibian declines through its effects on female body condition and survivorship. Oecologia, 151, 125–131. (doi: 10.1007/s00442-006-0558-1)

23. Arambourou H, Stoks R. 2015 Warmer winters modulate life history and energy storage but do not affect sensitivity to a widespread pesticide in an aquatic insect. Aquat. Toxicol., 167, 38–45. (doi: 10.1016/j.aquatox.2015.07.018)

24. Jørgensen CB. 1986 External and internal control of patterns of feeding, growth and gonadal function in a temperate zone anuran, the toad *Bufo bufo*. J. Zool., 210, 211–241. (doi: 10.1111/j.1469-7998.1986.tb03631.x)

25. Reading CJ, Clarke RT. 1995 The effects of density, rainfall and environmental temperature on body condition and fecundity in the common toad, *Bufo bufo*. Oecologia, 102, 453–459. (doi: 10.1007/BF00341357)

26. Meehl GA, Zwiers F, Evans J, Knutson T, Mearns L, Whetton P. 2000 Trends in extreme weather and climate events: issues related to modeling extremes in projections of future climate change. Bull. Am. Meteorol. Soc., 81, 427–436. (doi: 10.1175/1520-0477(2000)081<0427:TIEWAC>2.3.CO;2)

27. Ault TR, Henebry GM, De Beurs KM, Schwartz MD, Betancourt JL, Moore D. 2013 The false spring of 2012, earliest in North American record. Eos Trans. AGU, 94, 181–182. (doi: 10.1002/2013EO200001)

28. Ruthsatz K, Bartels F, Stützer D, Eterovick PC. 2022 Timing of parental breeding shapes sensitivity to nitrate pollution in the common frog *Rana temporaria*. J. Therm. Biol., 108, 103296. (doi: 10.1016/j.jtherbio.2022.103296)

29. Liu Q, Piao S, Janssens IA, Fu Y, Peng S, Lian XU, Ciais P, Myneni RB, Peñuelas J, Wang T. 2018 Extension of the growing season increases vegetation exposure to frost. Nat. Commun., 9, 426. (doi: 10.1038/s41467-017-02690-y)

30. Boutilier RG, Tattersall GJ, Donohoe PH. 2000 Metabolic consequences of behavioural hypothermia and oxygen detection in submerged overwintering frogs. Zool., 102, 111–119.

31. Sinsch U, Leskovar C. 2011 Does thermoregulatory behaviour of green toads (Bufo viridis) constrain geographical range in the west? A comparison with the performance of syntopic natterjacks (*Bufo calamita*). J. Therm. Biol., 36, 346–354. (doi: 10.1016/j.jtherbio.2011.06.012)

32. Costanzo JP, Lee Jr RE. 2013 Avoidance and tolerance of freezing in ectothermic vertebrates. J. Exp. Biol., 216, 1961–1967. (doi: 10.1242/jeb.070268)

33. Storey KB, Storey JM. 1988 Freeze tolerance in animals. Physiol. Rev., 68, 27–84. (doi: 10.1152/physrev.1988.68.1.27)

34. Storey JM, Storey KB. 1985 Triggering of cryoprotectant synthesis by the initiation of ice nucleation in the freeze tolerant frog, *Rana sylvatica*. J. Comp. Physiol. B, 156, 191–195. (doi: 10.1007/BF00695773)

35. Storey KB, Storey JM. 2017 Molecular physiology of freeze tolerance in vertebrates. Physiol. Rev., 97, 623–665. (doi: 10.1152/physrev.00016.2016)

36. do Amaral MCF, Frisbie J, Goldstein DL, Krane CM. 2018 The cryoprotectant system of Cope’s gray treefrog, *Dryophytes chrysoscelis*: responses to cold acclimation, freezing, and thawing. J. Comp. Physiol. B, 188, 611–621. (doi: 10.1007/s00360-018-1153-6)

37. Reading CJ. 1998 The effect of winter temperatures on the timing of breeding activity in the common toad *Bufo bufo*. Oecologia, 117, 469–475. (doi: 10.1007/s004420050682)

38. Kuhn J. 1994 Lebensgeschichte und Demographie von Erdkrötenweibchen Bufo bufo bufo (L.). Zeitschr. Feldherpet., 1, 3–87.

39. Orlofske SA, Hopkins WA. 2009 Energetics of metamorphic climax in the pickerel frog (*Lithobates palustris*). Comp. Biochem. Physiol. A: Mol. Integr. Physiol., 154, 191–196. (doi: 10.1016/j.cbpa.2009.06.001)

40. Peig J, Green AJ. 2009 New perspectives for estimating body condition from mass/length data: the scaled mass index as an alternative method. Oikos, 118, 1883–1891. (doi: 10.1111/j.1600-0706.2009.17643.x)

41. MacCracken JG, Stebbings JL. 2012 Test of a body condition index with amphibians. J. Herpetol., 46, 346–350. (doi: 10.1670/10-292)

42. Gabor CR, Bosch J, Fries JN, Davis DR. 2013 A non-invasive water-borne hormone assay for amphibians. Amphib.-Reptil., 34, 151–162. (doi: 10.1163/15685381-00002877)

43. Ruthsatz K, Rico-Millan R, Eterovick PC, Gomez-Mestre I. 2023 Exploring water-borne corticosterone collection as a non-invasive tool in amphibian conservation physiology: benefits, limitations and future perspectives. Conserv. Physiol., 11, coad070. (doi: 10.1093/conphys/coad070)

44. Burraco P, Valdés AE, Johansson F, Gomez-Mestre I. 2017 Physiological mechanisms of adaptive developmental plasticity in *Rana temporaria* island populations. BMC Evol. Biol., 17, 1–10. (doi: 10.1186/s12862-017-1004-1)

45. McCann SM, Kosmala G, Greenlees MJ, Shine R. 2018 Physiological plasticity in a successful invader: rapid acclimation to cold occurs only in cool-climate populations of cane toads (*Rhinella marina*). Conserv. Physiol., 6, cox072. (doi: 10.1093/conphys/cox072)

46. Taylor EN, Diele-Viegas LM, Gangloff EJ, Hall JM, Halpern B, Massey MD, Rödder D, Rollinson N, Spears S, Telemeco RS, et al. 2021 The thermal ecology and physiology of reptiles and amphibians: A user’s guide. J. Exp. Zool. A: Ecol. Integr. Physiol., 335, 13–44. (doi: 10.1002/jez.2396)

47. Wickham H. 2009 Elegant graphics for data analysis. Media, 35(211), 10–1007.

48. Yu D, Zhang Z, Glass K, Su J, DeMeo DL, Tantisira K, Weiss ST, Qiu W. 2019 New statistical methods for constructing robust differential correlation networks to characterize the interactions among microRNAs. Sci. Rep., 9, 3499. (doi: 10.1038/s41598-019-40167-8)

49. Chin WW. 1998 Commentary: Issues and opinion on structural equation modeling. MISQ, 22, vii–xvi. (doi: 10.2307/249674)

50. Grabherr MG, Haas BJ, Yassour M, Levin JZ, Thompson DA, Amit I, Adiconis X, Fan L, Raychowdhury R, Zeng Q, et al. 2011 Full-length transcriptome assembly from RNA-Seq data without a reference genome. Nat. Biotechnol., 29, Article 7. (doi: 10.1038/nbt.1883)

51. Haas BJ, Papanicolaou A, Yassour M, Grabherr M, Blood PD, Bowden J, Couger MB, Eccles D, Li B, Lieber M, et al. 2013 De novo transcript sequence reconstruction from RNA-seq using the Trinity platform for reference generation and analysis. Nat. Protoc., 8, Article 8. (doi: 10.1038/nprot.2013.084)

52. Langmead B, Salzberg SL. 2012 Fast gapped-read alignment with Bowtie 2. Nat. Methods, 9, 357–359. (doi: 10.1038/nmeth.1923)

53. Patro R, Duggal G, Love MI, Irizarry RA, Kingsford C. 2017 Salmon provides fast and bias-aware quantification of transcript expression. Nat. Methods, 14, Article 4. 10.1038/nmeth.4197

54. Love MI, Huber W, Anders S. 2014 Moderated estimation of fold change and dispersion for RNA-seq data with DESeq2. Genome Biol., 15, 550. (doi: 10.1186/s13059-014-0550-8)

55. Kolde R. 2019 pheatmap: Pretty Heatmaps. R Pack. Version, 1.0. 12.

56. Bryant DM, Johnson K, DiTommaso T, Tickle T, Couger MB, Payzin-Dogru D, Lee TJ, Leigh ND, Kuo TH, Davis FG, et al. 2017 A Tissue-Mapped Axolotl De Novo Transcriptome Enables Identification of Limb Regeneration Factors. Cell Rep., 18, 762–776. (10.1016/j.celrep.2016.12.063)

57. Camacho C, Coulouris G, Avagyan V, Ma N, Papadopoulos J, Bealer K, Madden TL. 2009 BLAST+: Architecture and applications. BMC Bioinform., 10, 421. (doi: 10.1186/1471-2105-10-421)

58. UniProt Consortium. 2019 UniProt: a worldwide hub of protein knowledge. Nucleic acids research, Nucleic Acids Res., 47, D506–D515. (doi: 10.1093/nar/gky1049)

59. Thomas PD, Ebert D, Muruganujan A, Mushayahama T, Albou LP, Mi H. 2022 PANTHER: Making genome-scale phylogenetics accessible to all. Protein Sci., 31, 8–22. (doi: 10.1002/pro.4218)

60. Angilletta Jr MJ, Niewiarowski PH, Navas CA. 2002 The evolution of thermal physiology in ectotherms. J. Therm. Biol., 27, 249–268. (doi: 10.1016/S0306-4565(01)00094-8)

61. Glennemeier KA, Denver RJ. 2002 Small changes in whole-body corticosterone content affect larval *Rana pipiens* fitness components. Gen. Comp. Endocr., 127, 16–25. (doi: 10.1016/S0016-6480(02)00015-1)

62. Sapolsky RM, Romero LM, Munck AU. 2000 How do glucocorticoids influence stress responses? Integrating permissive, suppressive, stimulatory, and preparative actions. Endocr. Rev., 21, 55–89. (doi: 10.1210/edrv.21.1.0389)

63. Crespi EJ, Williams TD, Jessop TS, Delehanty B. 2013 Life history and the ecology of stress: how do glucocorticoid hormones influence life-history variation in animals?. Funct. Ecol., 27, 93–106. (doi: 10.1111/1365-2435.12009)

64. Romero LM, Beattie UK. 2022 Common myths of glucocorticoid function in ecology and conservation. J. Exp. Zool. A: Ecol.Integr. Physiol., 337, 7–14. (doi: 10.1002/jez.2459)

65. Kirschman LJ, McCue MD, Boyles JG, Warne RW. 2017 Exogenous stress hormones alter energetic and nutrient costs of development and metamorphosis. J. Exp. Zool., 220, 3391–3397. (doi: 10.1242/jeb.164830)

66. Jimeno B, Verhulst S. 2023 Meta-analysis reveals glucocorticoid levels reflect variation in metabolic rate, not ‘stress’. Elife, 12, RP88205. (doi: 10.7554/eLife.88205.3)

67. Wilsterman K, Ballinger MA, Williams CM. 2021 A unifying, eco-physiological framework for animal dormancy. Funct. Ecol., 35, 11–31. (doi: 10.1111/1365-2435.13718)

68. Rollins-Smith LA, Woodhams DC. 2012 Ecoimmunology. Demas G., Nelson R. (eds.), Oxford University Press, New York, 92–143

69. Benard MF. 2015 Warmer winters reduce frog fecundity and shift breeding phenology, which consequently alters larval development and metamorphic timing. Glob. Chang. Biol., 21, 1058–1065. (doi: 10.1111/gcb.12720)

70. Rollins-Smith LA. 2017 Amphibian immunity–stress, disease, and climate change. Dev. Comp. Immunol., 66, 111–119. (doi: 10.1016/j.dci.2016.07.002)

71. Lavy D, Nedved O, Verhoef HA. 1997 Effects of starvation on body composition and cold tolerance in the collembolan Orchesella cincta and the isopod Porcellio scaber. J. insect physiol., 43, 973–978. (doi: 10.1016/S0022-1910(97)00011-5)

72. Pörtner HO, Van Dijk PLM, Hardewig I, Sommer A. 2000 Levels of metabolic cold adaptation: tradeoffs in eurythermal and stenothermal ectotherms. Antarctic Ecosystems: models for wider ecological understanding. Davison W., Howard Williams C. (eds.), Christchurch New Zealand, 109–122.

73. Sobek-Swant S, Crosthwaite JC, Lyons DB, Sinclair BJ. 2012 Could phenotypic plasticity limit an invasive species? Incomplete reversibility of mid-winter deacclimation in emerald ash borer. Biol. Invasions, 14, 115–125. (doi: 10.1007/s10530-011-9988-8)

74. Sinclair BJ, Stinziano JR, Williams CM, MacMillan HA, Marshall KE, Storey KB. 2013 Real-time measurement of metabolic rate during freezing and thawing of the wood frog, *Rana sylvatica*: implications for overwinter energy use. J. Exp. Biol., 216, 292–302. (doi: 10.1242/jeb.076331)

75. Bale JS, Hayward SAL. 2010 Insect overwintering in a changing climate. J. Exp. Biol., 213, 980–994. (doi: 10.1242/jeb.037911)

76. Storey KB. 1987 Organ-specific metabolism during freezing and thawing in a freeze-tolerant frog. Am. J. Physiol. Regul. Integr. Comp. Physiol., 253, R292–R297.

77. Just PA, Charawi S, Denis RG, Savall M, Traore M, Foretz M, … & Perret C. 2020 Lkb1 suppresses amino acid-driven gluconeogenesis in the liver. Nat. Comm., 11, 6127. (doi:10.1038/s41467-020-19490-6)

78. Denzel MS, Antebi A. 2015 Hexosamine pathway and (ER) protein quality control. Curr. Opin. Cell Biol., 33, 14–18. (doi: 10.1016/j.ceb.2014.10.001)

79. Nakajima A, Yamaguchi R, Sasazaki M, Ishihara A, Yamauchi K. 2023 Adult male *Xenopus laevis* can tolerate months of fasting by catabolizing carbohydrates and lipids. J. Comp. Physiol. B, 193, 227–238. (doi: 10.1007/s00360-023-01478-5)

80. de Amaral M, Von Dentz MC, Simões LAR, Vogt É, Heiermann D, Fischer P, Colombo P, Kucharski LC. 2023 Metabolic changes in the subtropical frog *Boana pulchella* during experimental cooling and recovery conditions. J. Therm. Biol., 117, 103705. (doi: 10.1016/j.jtherbio.2023.103705)

81. Schmidt R, Zummach C, Sinai N, Sabino-Pinto J, Künzel S, Dausmann KH, Ruthsatz K. 2024 Physiological responses to a changing winter climate in an early spring-breeding amphibian. bioRxiv, 2024-01. (doi: 10.1101/2024.01.03.574103)

82. Gittins SP, Parker AG, Slater FM. 1980 Population characteristics of the common toad (*Bufo bufo*) visiting a breeding site in mid-Wales. J. Anim. Ecol., 161–173. (doi: 10.2307/4281)

83. Sinsch U. 1988 Seasonal changes in the migratory behaviour of the toad *Bufo bufo*: direction and magnitude of movements. Oecologia, 76, 390–398. (doi: 10.1007/BF00377034)

84. Podhajský L, Gvoždík L. 2016 Variation in winter metabolic reduction between sympatric amphibians. Comp. Biochem. Physiol. A: Mol. Integr. Physiol., 201, 110–114. (doi: 10.1016/j.cbpa.2016.07.003)

